# Diet-induced rewiring of the Wnt gene regulatory network connects aberrant splicing to fatty liver and liver cancer in DIAMOND mice

**DOI:** 10.1101/2023.02.09.527844

**Authors:** Ana López-Pérez, Silvia Remeseiro, Andreas Hörnblad

## Abstract

**Background & Aims:** This study aimed to provide a comprehensive understanding of the regulatory and transcriptional landscape in liver tumours from DIAMOND mice, a mouse model that mimics human hepatocellular carcinoma (HCC) in the context of metabolic associated fatty liver disease (MAFLD).

**Methods:** RNA-sequencing and ChIP-sequencing were used to study the gene expression and regulatory changes in DIAMOND liver tumours. RNA *in situ* hybridisation splice variant analysis was used to study β-catenin exon 3 exclusion in tumours at cellular resolution. Sequencing data on β-catenin exon 3 splicing in DIAMOND tumours was compared to data from human patients and cell lines.

**Results:** The study found an increase in Wnt/β-catenin-signalling accompanied by rewiring of the Wnt/β-catenin regulatory network in DIAMOND tumours. Changes include switching in the expression of the canonical TCF/LEF downstream effectors and associated chromatin remodelling. In addition, a large subset of DIAMOND tumours showed aberrant splicing of β-catenin, which generate an mRNA isoform that encodes an oncogenic protein. Similar splicing events were found in a fraction of human HCC and hepatoblastoma samples.

**Conclusions:** This study provides evidence that western diet induces aberrant genome-wide splicing in DIAMOND livers, and in particular of the β-catenin gene in a subset of DIAMOND liver tumours. This mechanism is distinct from previously reported activation of β-catenin in HCC and mouse models, since it is independent on mutations in the locus. Our data suggests that metabolic input modulates gene regulatory network responses to active Wnt-signalling, which will be an important consideration also in the human setting.

**Lay summary:** Sequencing data generated in this study highlights the effect of diet in modulating oncogenic gene expression and underscores an alternative mutation-independent mechanism leading to constitutive activation of β-catenin, a well-known driver of liver cancer.

## Introduction

Liver cancer is among the leading causes for cancer-related death worldwide and the projection is that almost 1,5 million people will be affected by 2040 [1]. Importantly, many of the newly diagnosed cases in developed countries are associated with obesity, metabolic dysfunction and associated complications, including Metabolic Associated Fatty Liver Disease (MAFLD). Obesity is predicted to become the leading cause for liver cancer in western countries, and fatty-liver-associated cancer will thus contribute increasingly to liver-cancer mortality worldwide [2,3].

Molecular classification of HCC broadly defines two subclasses: proliferative and non-proliferative HCC. The proliferative class is characterized by gene signatures of poor prognosis, often carry TP53 mutations and has a worse clinical outcome, while the non-proliferative class frequently carries β-catenin mutations and is associated with a better clinical outcome [4]. *TRP53* and *CTNNB1* are also some of the most frequently mutated genes in HCC (each in ~30% of cases), only mutations in the *TERT* promoter are more frequent (~60%) [4]. Importantly, *CTNNB1* mutations are most often found in the absence of Hepatitis B virus infection and have been associated with liver cancer in the context of metabolic syndrome [5]. These coding mutations disrupt the β-catenin degradation domain encoded in the exon 3 of *CTNNB1*, either via point mutations or larger in-frame deletions that remove key residues that upon phosphorylation target the protein for degradation [5]. This leads to stabilization of β-catenin in the cytosol and subsequent translocation into the nucleus [6]. In the nucleus β-catenin interacts with LEF/TCF transcription factors to regulate gene expression. Recent reports have demonstrated that similar activation of β-catenin, either via engineered deletions or CRISPR targeting of *Ctnnb1* exon 3 in the liver, is sufficient to generate tumours in mice [7,8].

Recent evidence highlights how mutations in non-coding regulatory elements or disruptions of 3D chromatin architecture can lead to aberrant gene expression in cancer [9–13]. It is also well known that dysfunction of other regulatory mechanisms such as alternative splicing contribute importantly to cancer aetiology [14] including HCC [15–17]. Still, a thorough understanding on the role of the regulatory genome or altered splicing patterns in the aetiology of MAFLD-associated HCC is lacking.

To better characterize the molecular underpinnings of liver cancer aetiology in the context of metabolic syndrome and fatty liver disease, various mouse models have been developed in recent years [18–22]. Among them, the DIAMOND mouse model mimics several aspects of MAFLD-driven liver cancer. This mouse model is generated upon feeding western diet (WD) to an isogenic strain isolated from an interbreeding between C57Bl/6J and Sv129/vImJ, and that in response develop MAFLD and liver tumours with an incidence of ~90% [19].

Herein, we present an in-depth analysis of the transcriptional and regulatory landscape of DIAMOND liver tumours and fatty liver tissue, in comparison to healthy litter mate controls fed regular diet. The study describes extensive re-wiring of the Wnt gene regulatory network in DIAMOND mice and demonstrates that alternative splicing is a key feature of MAFLD and HCC in this model. More specifically, we identified diet-induced aberrant splicing, independent on locus-specific somatic mutations, as one molecular driver underlying oncogenic β-catenin-signalling in a large subset of these mouse liver tumours. Similar aberrant splicing of β-catenin in human liver cancer samples were also observed, highlighting the potential importance also in the development of human liver cancer.

## Results

### DIAMOND mice develop HCC in the context of obesity and insulin resistance

To investigate in depth the genome-wide transcriptional and epigenetic changes that occur during development of MAFLD and obesity-related hepatocellular carcinoma (HCC), we performed RNA-sequencing and Chromatin ImmunoPrecipitation-sequencing (ChIP-seq) of liver tumours (HCC) and liver control tissue (FL) from DIAMOND mice, as well as liver tissue from healthy control mice fed a regular diet (RD) (Figure 1A). The feeding regimens were introduced at 8 weeks of age and mice were sacrificed at approximately one year after the start of the diet. In accordance with previous findings [19], DIAMOND mice developed obesity, insulin resistance and dyslipidemia (Figure S1-S2). There were no significant changes in fasting blood glucose or plasma insulin levels at the start of the diet, nor after 42 weeks of diet (Figure S1C). However, Insulin Tolerance Test (ITT) at T=16w weeks as well as Glucose Tolerance (GTT) and Glucose Stimulated Insulin Secretion (GSIS) tests at T=19w demonstrated significant insulin resistance in the DIAMOND mice compared to the controls (Figure S1D-F).

**Figure 1.**
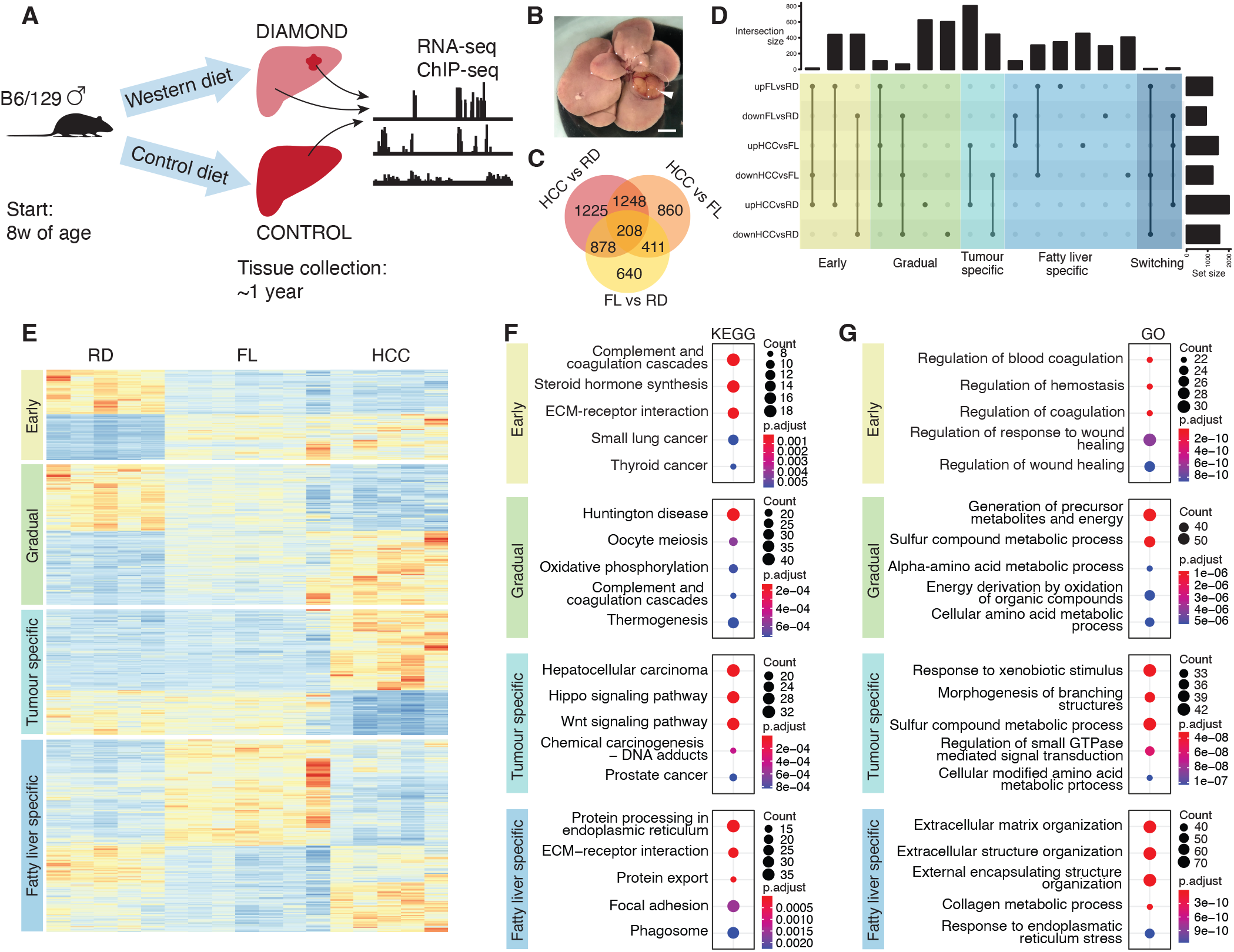
Gene expression changes associated with HCC development in DIAMOND mice. **A**) Schematic workflow. **B**) Photomicrograph of DIAMOND liver with tumour. Scale bar corresponds to 5mm. **C**) Venn diagram of differentially expressed genes (DEGs) in pairwise comparisons between RD, FL and HCC. **D**) Upset plot of differentially expressed genes categorised as early, gradual, tumor-specific, fatty liver specific and switching. **E**) Heatmap of gene expression levels for early, gradual, tumor-specific and fatty liver-specific genes. Enrichment of KEGG (**F**) and GO (**G**) terms for each of the defined categories.

Using the Vetscan 2 “mammalian liver profile” system to assess liver function at T=38w, we observed increased blood levels of various factors in DIAMOND mice compared to healthy controls, including alkaline phosphatase, aspartate transaminase, bile acids, albumin, urea nitrogen and cholesterol (Figure S2A). Also, at the end-point liver triglyceride and cholesterol levels were significantly increased in the DIAMOND mice (Figure S2B). These data indicated both significant liver damage and dyslipidemia in DIAMOND mice as compared to littermate controls. Only 25% of the control livers (2 out of 8) carried tumours, while ~89% (16 out of 18) of the DIAMOND livers had one or several tumours at the endpoint of the experiment. Taken together, these data corroborate previously published data [19] on glucose and lipid homeostasis as well as tumour incidence in these mice, and demonstrate the robustness of the phenotypes reported for this model, also in our hands.

### Transition to malignancy is associated with increased activation of Wnt-signalling

RNA-seq analysis demonstrated that 2727 genes were differentially expressed in HCC tumours compared to FL samples, and 3559 genes as compared to RD samples. Out of these, 1248 genes were differentially expressed in both comparisons (Figure 1C). To better understand these transcriptional changes that take place in the development of MAFLD and associated HCC in these mice, and to identify gene expression changes that are likely associated with the malignant phenotype, we grouped the genes into 5 distinct categories based on their expression profiles (Figure 1D-E): early (892), gradual (1396), tumour-specific (1248), fatty liver-specific (1911), and switching (23) genes. We defined early genes as genes where expression changes are evident already in FL tissue and maintained also in HCC. Gradual refers to gradual changes from RD to FL to HCC. Tumour-specific and fatty liver-specific were only differentially expressed in HCC or FL tissue, respectively. Switching genes were defined as dysregulated in opposite directions in FL and HCC compared to RD samples (i. e. up in FL, down in HCC, or down in FL, up in HCC). Next, we performed enrichment analysis of gene ontology terms (GO) and KEGG pathways for these 5 gene categories. The most enriched KEGG and GO terms for early genes were related to coagulation cascades and wound healing (Figure 1F-G, Table S1). Although this was also enriched for gradual genes, the most highly enriched terms were related to metabolism, metabolic function, and cell cycle (Figure 1F-G, Table S1). This was further reflected by enrichment of terms related to mitochondria, cellular respiration, and cell cycle in the GO cellular compartment (Table S1). Of note, although not among the top 5 most highly enriched KEGG terms, NAFLD (Non-Alcoholic Fatty Liver Disease, previous nomenclature for MAFLD) was also enriched (Table S1). Analysis of the tumour-specific genes revealed hepatocellular carcinoma, hippo- and Wnt-signalling pathways among the top three enriched KEGG terms, but also identified GO terms related to morphogenesis and developmental processes, including Wnt-signalling (Figure 1F-G, Table S1). This confirms DIAMOND mice as a relevant model for human HCC, especially in the context of aberrant Wnt-activation. Although there is a large group of genes that we defined as gradual, most dysregulated genes (1911 out of 2137) in non-tumorous fatty liver tissue compared to RD are fatty liver-specific. These FL-specific genes appear to be largely associated with fibrotic processes as they were enriched for terms related to extracellular matrix organization, collagen, and protein processing (Figure 1F-G, Table S1). This is in concordance with the different morphology of the tumours and fatty liver tissue, where fibrosis was only present in the latter (Figure S3). Lastly, the very few members among the switching genes were enriched for a diverse set of terms including tryptophan, xenobiotic and retinol metabolism, as well as steroid hormone biosynthesis (Table S1). Taken together, these analyses demonstrate extensive differences in the transcriptional landscape of RD livers compared to FL and HCC, but also between FL and HCC and highlight extensive transcriptional changes in the Wnt-signalling pathway as one main driver of tumorigenesis in this model.

### Gradual reorganisation of the chromatin landscape in DIAMOND liver tumours

Spatial and temporal specificity of gene expression rely, among others, on non-coding regulatory sequences in the genome such as enhancers. These can be located far (up to megabases) from their target genes in the linear genome and their regulatory activity is intimately linked to their epigenetic status [23,24]. To chart the gene regulatory changes that take place during HCC development, we performed ChIP-seq for the active enhancer (H3K27ac) and repressive (H3K27me3) chromatin marks in tumour (HCC, n=3) and non-tumour (FL, n=2) liver tissue from DIAMOND mice, as well as in healthy control livers (RD, n=3). Genomic annotations of H3K27ac regions were similar in RD, FL and HCC; however, H3K27me3 peaks in RD livers were more frequently located close to transcriptional start sites (<5kb from TSS) as compared to FL and HCC peaks (Figure 2A). Importantly, differential binding analysis revealed significant redistribution of active and repressive chromatin marks in the three conditions (Figure 2B-D).

**Figure 2.**
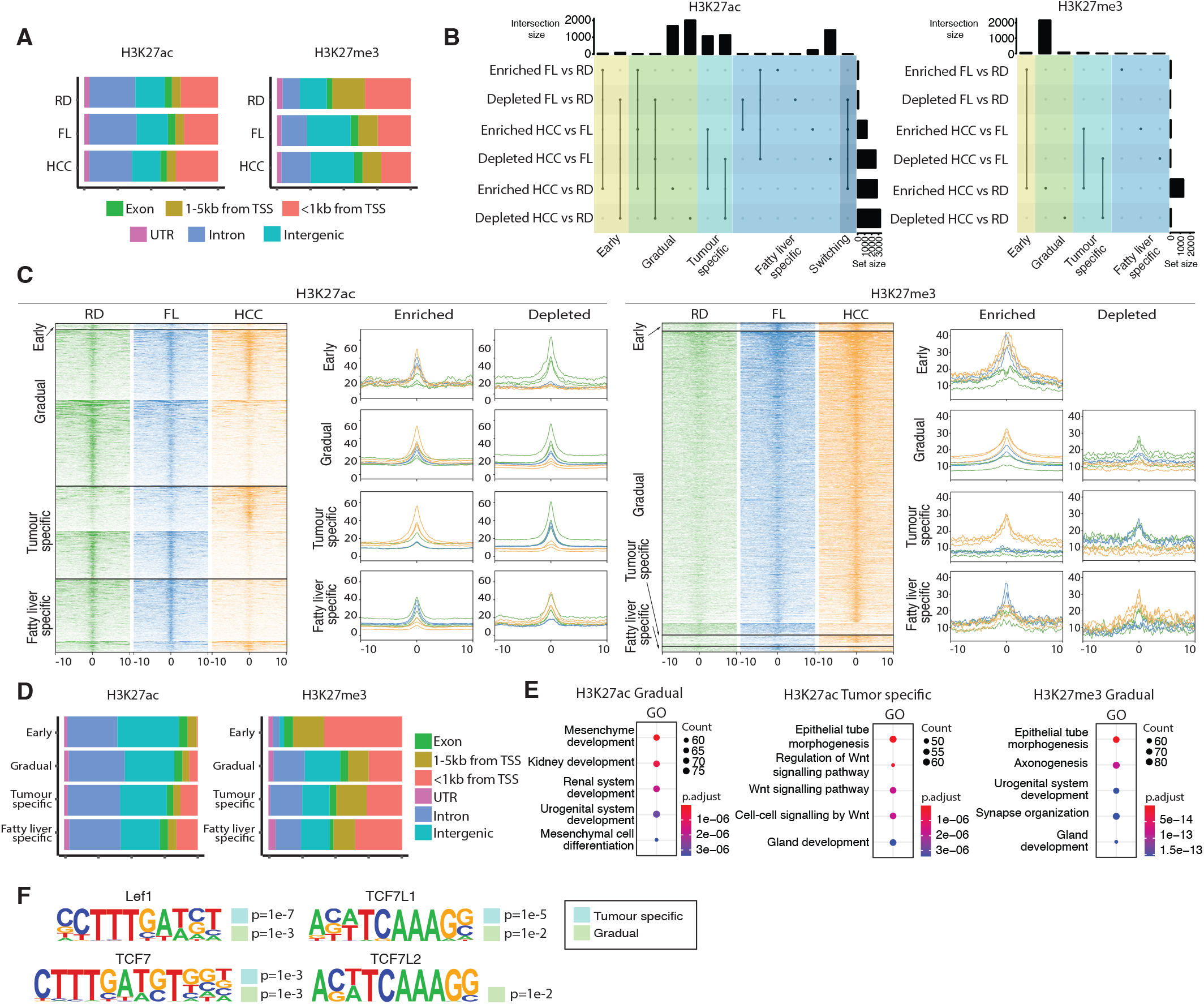
Tumour-associated active regulatory regions integrate transduction of Wnt-signalling. **A**) Genomic distribution of H3K27ac and H3K27me3 chromatin marks in RD, FL and HCC tissue. **B**) Upset plot of differentially binding regions for H3K27ac and H3K27me3 in pairwise comparisions between RD, FL and HCC, and categorised as early, gradual, tumor-specific, fatty liver specific and switching regions. **C**) Heat map and density plots depicting read distribution ±10kb around differential binding regions of H3k27ac and H3K27me3 for the defined categories. **D**) Genomic distribution of H3K27ac (left) and H3K27me3 (right) for early, gradual, tumour-specific and fatty-liver specific regions. **E**) Enrichment of GO terms for gradual and tumour-specific H3K27ac regions, and for gradual H3K27me3 regions. **F**) Tcf/Lef motifs present in tumour-specific (marked by light blue box) and gradual regions (light green box) enriched for H3K27ac.

To better understand these regulatory changes that take place, and similar to the RNA-seq analysis, we defined differential regions as early, gradual, tumour-specific, fatty liver-specific, and switching (Figure 2B-C, Table S2, Table S3). This revealed that most of the redistribution of H3K27ac occurred gradually. In fact, these changes made up ~48% of the total number of differential regions, while tumour-specific and fatty liver-specific regions constituted ~28% and 22% of the differential regions, respectively. Less than 2% of the peaks were associated with early or switching regions. In gradual and tumour-specific regions loss or gain of H3K27ac occurred with similar frequency (54/46% depletion/enrichment, and 51/49% depletion/enrichment, respectively) while fatty-liver specific regions were strongly biased towards enrichment (~85%). The by far most frequent change in repressive H3K27me3 was the gain of gradual regions (~93% of all differential binding events).

Among the differential regions, H3K27ac peaks were mostly located in introns and intergenic with a small proportion (<13%) located to promoter regions (<1kb from TSS) (Figure 2D). In contrast, differential H3K27me3-regions were more frequently located within 5kb from the nearest gene (~50%, Figure 2D).

Thus, our study shows a gradual remodelling of the gene regulatory landscape in obesity-related HCC in this model. This data suggests that the chromatin environment formed in the liver under conditions of overnutrition is an important factor contributing to a pro-tumorigenic niche that can trigger HCC.

### Tumour-associated active regulatory regions integrate transduction of Wnt-signalling

To further investigate how redistribution of chromatin marks related to the molecular underpinnings of tumorigenesis, we performed GO analysis for the genes closest to the differential regions for both chromatin marks (Table S2-3). The most highly enriched GO terms for genes in close proximity to tumour-specific H3K27ac regions were associated to Wnt-signalling. So were the genes near gradual regions, albeit to a lesser extent (Figure 2E). The gradual regions were most significantly associated with genes related to development and developmental processes (Figure 2E). For H3K27me3, a large majority of the differential regions were gradual regions, and similarly to the gradual H3K27ac regions, these were associated to genes involved in developmental processes, and in particular axonogenesis and synapse organisation (Figure 2E, Table S3). This is interesting in view of recent reports on nerve degeneration in MAFLD [25] and the emerging data on the role of the neuron-tumour crosstalk in malignancies in general [26]. Importantly, also for these gradual regions, Wnt-related GO terms were enriched.

Next, we investigated the putative TF binding motifs in differentially binding regions, assessing enriched and depleted regions separately. Importantly, the most significantly enriched TF motif in tumour-specific H3K27ac regions was LEF1, while TCF7 and TCF7L1 motifs were also highly enriched (Figure 2F). These together with TCF7L2 are the classical effectors of the canonical Wnt-pathway. Motifs for all four of these were also present in gradually enriched regions (Figure 2F). In contrast, TCF/LEF motifs were not enriched in H3K27ac-depleted regions in tumours nor in gradually depleted H3K27ac regions. Also, of the differential H3K27me3 peaks, only the fatty liver-specifically enriched regions were associated with the canonical Wnt-signalling via enrichment of TCF7L2 motifs. The gradually repressed regions, which constituted the lion part of differential H3K27me3 binding, were most highly enriched for IRF8 binding motifs. These motifs were also present in the gradually depleted and tumour-specific depleted H3K27ac. This is interesting, as IRF8 has been proposed to suppress HCC progression [27] and in the context of leukemia, depletion of IRF8 in combination with activated β-catenin enhances malignant gene expression profile [28]. Taken together, these data indicate that canonical Wnt-signalling is the major direct activator of aberrant gene expression program in DIAMOND tumours.

### Rewiring of the Wnt/β-catenin gene regulatory network accompanies DIAMOND liver tumorigenesis

Given the enrichment of Wnt/β-catenin GO terms and TCF/LEF transcription factor binding motifs in tumour-associated DEGs and active chromatin, respectively, we explored in detail the RNA-seq gene expression data for individual core components of the canonical Wnt/β-catenin-pathway. A tumour-specific upregulation of *Lef1* and *Tcf7* genes was observed, as well as downregulation of *Tcf7l1*, while *Tcf7l2* expression was unaltered (Figure 3A). Upregulation of *Lef1* expression was accompanied by redistribution of chromatin marks, including gain of active marks in regions both proximal and distal to the gene promoter (Figure 3B). Furthermore, *Ctnnb1* expression was increased in tumours compared to both FL and RD livers and among the components of the β-catenin destruction complex, the known targets *Axin1* and *Axin2* were significantly upregulated, while *Apc, Gsk3a* and *Gsk3b* were not differentially expressed. Increased *Axin2* expression was also associated with tumour-specific H3K27ac enrichment in several regions surrounding the gene. In line with increased activation of Wnt-signalling, the downstream targets *Lgr5* and *Myc* were also significantly upregulated. Thus, DIAMOND tumorigenesis is not only accompanied by upregulation of Wnt-signalling and associated chromatin remodelling, but also by a rewiring in the downstream effectors, which is likely to also affect the set of downstream target genes and contribute to a malignant gene regulatory program.

**Figure 3.**
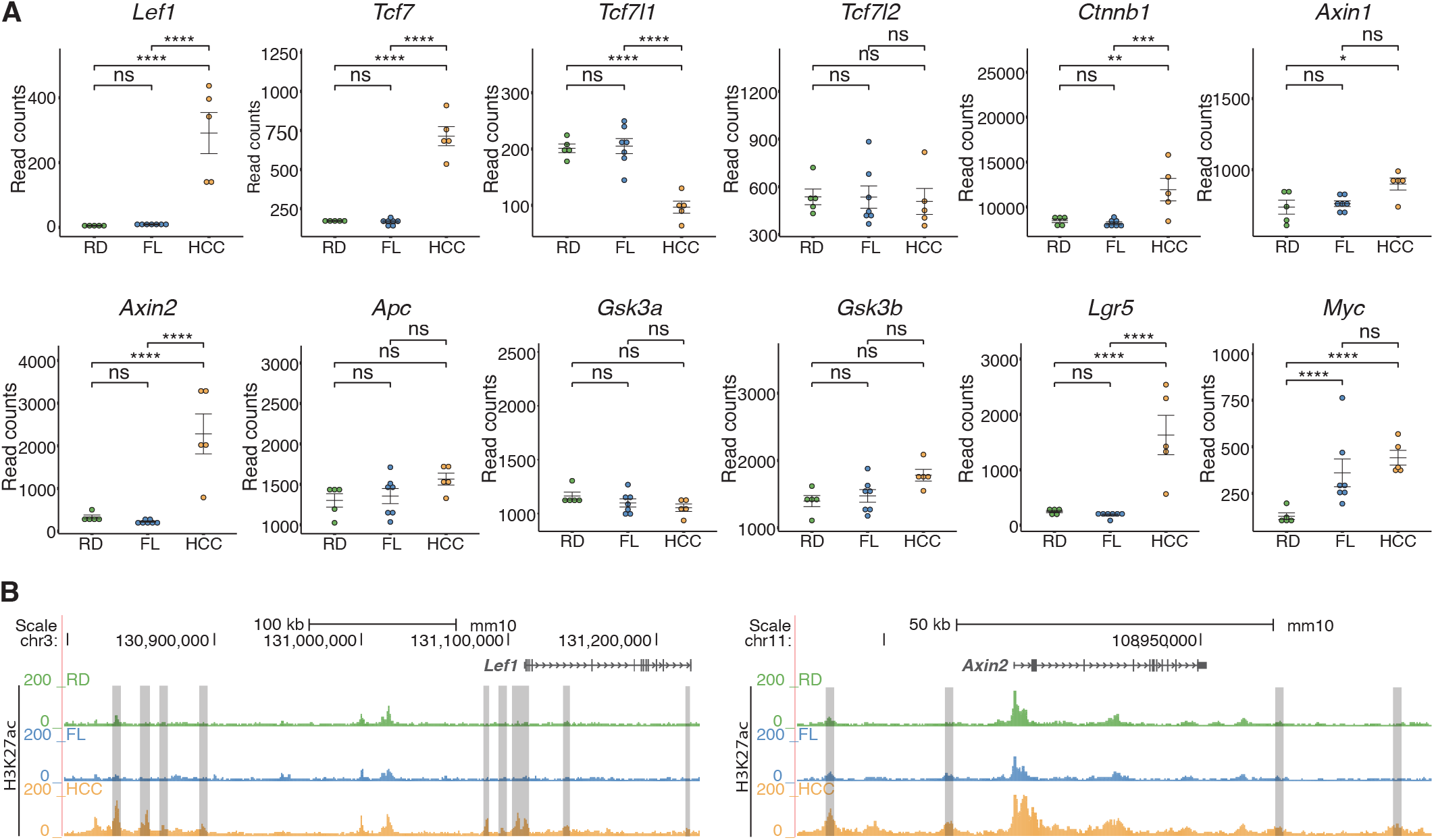
Rewiring of the Wnt-Ctnnb1 gene regulatory network accompanies DIAMOND liver tumorigenesis. **A**) Normalized read counts for core components of the canonical Wnt-Ctnnb1-pathway in RD, FL and HCC tissue. A clear switch in the expression of TF effectors can be seen in tumour tissue, where *Lef1* and *Tcf7* are upregulated while *Tcf7l1* levels are down. Individual data points, mean ± SEM are indicated. *p.adj.< 0.05, **p.adj.<0.01, ***p.adj.<0.001, ****p.adj.<0.0001, ns = not significant. **B**) Redistribution of H3K27ac in the *Lef1* and *Axin2* loci. Grey bars indicate H3K27ac regions that are gained specifically in tumours.

### Western diet induces distinct alternative splicing patterns in DIAMOND livers and links aberrant splicing in tumours to Wnt-signalling

Apart from the genome-wide regulatory and gene expression changes that occur in liver tumours in DIAMOND mice, we also investigated the general splicing patterns in the HCC tumours, FL and RD control liver tissue. Although it is known that cancer cells often display aberrant splicing, only few studies [29–32] have addressed the role of splicing in the context of MAFLD. Interestingly, our analysis revealed a significant global increase in the number of alternative splice events (ASE), not only in HCC, but also in FL tissue (Figure 4A). This quantitative difference reflects an increased number of ASEs per spliced gene as well as a higher number of genes that are subjected to alternative splicing (Figure 4B,C), meaning that it is not merely a consequence of differences in transcript abundance. Of note, 60 genes involved in mRNA-splicing (out of 296, GO:000398, “mRNA splicing, via spliceosome”) are also differentially expressed in our RNA-seq data. Unsupervised clustering based on these genes accurately group the samples into RD, FL, and HCC, where the FL cluster is more similar to RD than to tumours (Figure 4D). Interestingly, analysis of published RNA-seq data from human patients also demonstrates that quantitative differences in ASEs are associated with human MAFLD (Figure 4E).

**Figure 4.**
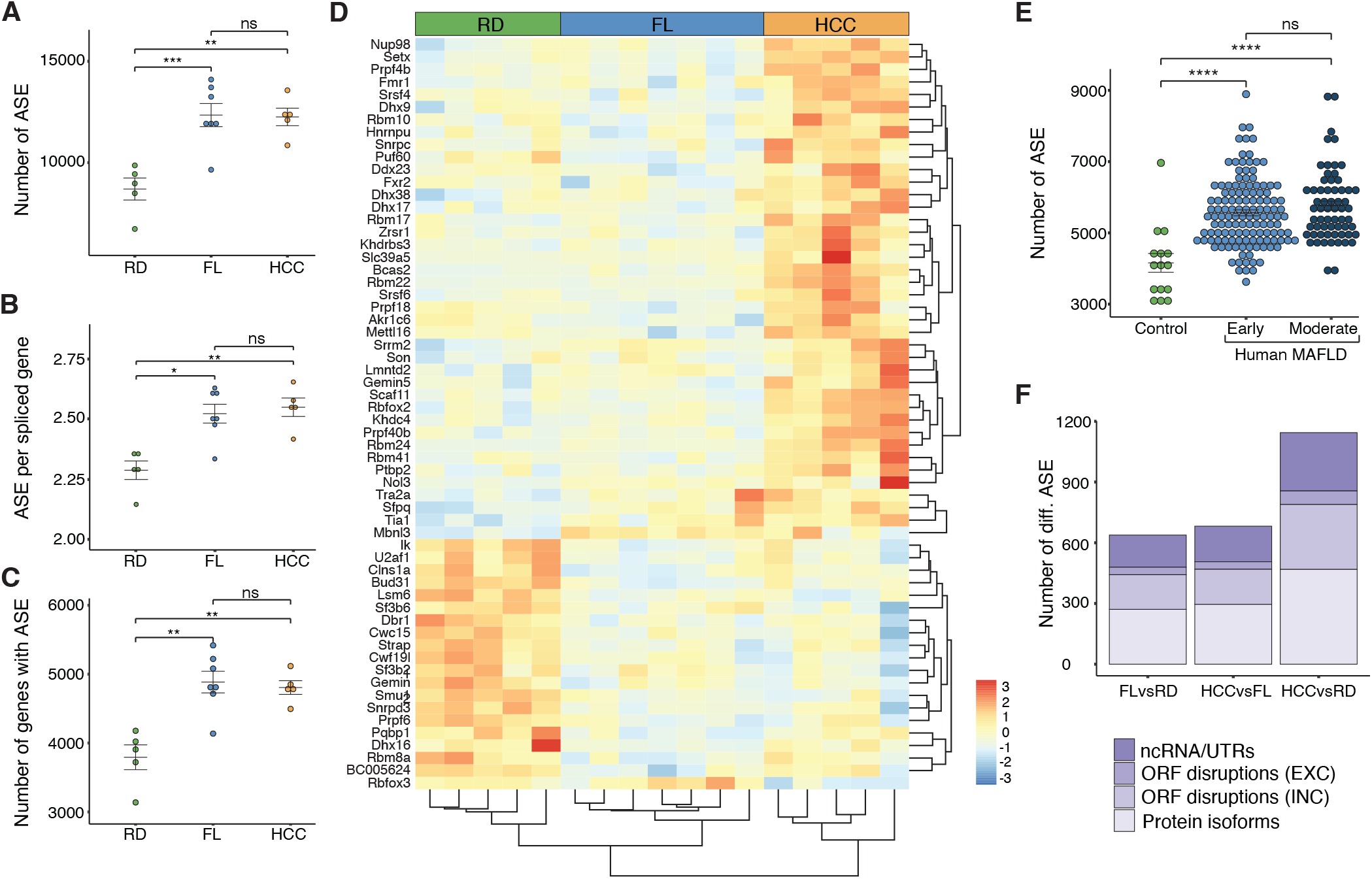
Changes in alternative splicing and altered expression of splice factors accompanies FL and HCC in DIAMOND mice. **A**) Total number of ASE (Alternative Splice Events) per sample for RD livers, FL and HCC tissue in DIAMOND mice. **B)** Frequency of ASE per gene and sample, and **C)**total number of genes with ASE per sample, for same mice as in **(A)**. Individual data points, mean ± SEM are indicated. *p < 0.05, **p<0.01, ***p<0.001 (Student’s t-test) **D)** Unsupervised clustering of RD, FL and HCC tissue based on 60 differentially expressed splice factors. Scale from red to blue indicates fold change over average read count for each row. **E)** Number of alternative splice events in human MAFLD livers compared to controls. Early and moderate indicate severity of MAFLD {Govaere:2020stm}. Individual data points, mean ± SEM are indicated. ****p<0.0001, ns = not significant (Student’s t-test) **F)** Average number of differential ASE in pairwise comparisions between RD, FL and HCC. Legend indicates ASEs leading to changes in ncRNA or UTRs, ORF disruptions due to exon exclusion (EXC), ORF disruptions due to intron inclusion (INC) or alternative protein isoforms.

To further dissect the observed changes in splicing patterns for our samples, a differential ASEs analysis was performed. Although the total number of ASEs were similar in FL and HCC tissues, the number of differential ASEs compared to RD was larger in HCC than in FL, indicating qualitative differences in the actual splicing events. Still, the proportion of different types of ASEs were similar for the differential ASEs in all comparisons (Figure 4F). In both FL and HCC differential ASEs were enriched for genes related to splicing and mRNA processing as compared to RD. Strikingly, also genes associated with Wnt-signalling and the β-catenin-TCF-complex were enriched (Table S4).

Our data shows that western diet in this model induces extensive gene regulatory changes at the level of splicing that may be an important part of the general disease aetiology. It also suggests that rewiring of the Wnt gene regulatory network in this context involves both transcriptional and co-transcriptional mechanisms, specifically connecting aberrant splicing to Wnt-signalling.

### Constitutive activation of β-catenin by alternative splicing contributes to Wnt/β-catenin -driven transcriptional activation in DIAMOND tumours

The enrichment of Wnt-signalling among the biological processes identified in our differential ASE analysis prompted us to further investigate specific genes in this pathway in more detail. The *Ctnnb1* gene was of particular interest due to its high frequency of mutations in liver cancer and its crucial role as the key effector in canonical Wnt-signalling. Strikingly, we identified a *Ctnnb1* transcript isoform excluding exon 3 in 60% (3 out of 5) of the sequenced DIAMOND tumours, where 25% of the *Ctnnb1* reads mapped to this region displayed exon 3 skipping (Figure 5A). Further RT-PCR analysis followed by Sanger sequencing revealed that in total 8 out of 18 (~44%) tumours presented *Ctnnb1* transcripts with exon 3 exclusion (Figure S4A-B). Exclusion of exon 3 was not detected in RD or FL tissue. No somatic mutations were detected in the *Ctnnb1* exon 3 region in 6 out of these 8 tumours displaying exon 3 exclusion (Figure S4C). This is relevant since it suggests that aberrant splicing underlies *Ctnnb1* exon 3 exclusion in this model, and in contrast to previous reports, it is not dependent on genomic mutations (e.g. alterations of regulatory splice sequences) in the locus, but it is rather induced by other mechanisms such as alterations in the splicing machinery.

**Figure 5.**
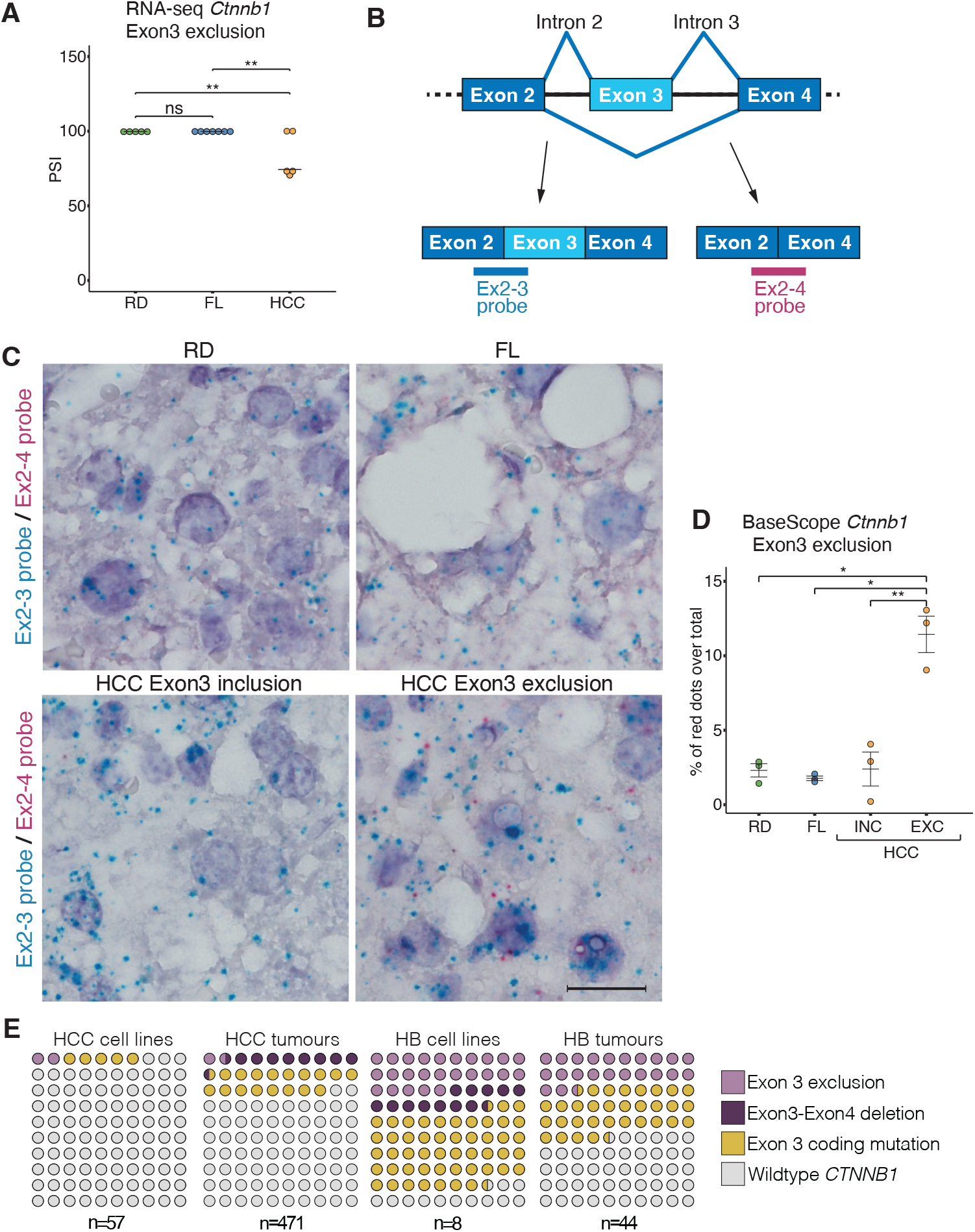
WD induces exclusion of Ctnnb1 in DIAMOND tumours. **A**) Exon-inclusion ratio (percent spliced in, PSI) for *Ctnnb1* exon3 in RD, FL and HCC RNA-seq samples. Individual data points and vast-tools estimated PSI are indicated. **p<0.01, ns = not significant. **B)** Schematic of exon 3 inclusion and exclusion transcripts and site of BaseScope probe annealing. **C)** Photomicrograph of representative sections from RD (upper left), FL (upper right), and HCC *Ctnnb1* exon 3 inclusion (lower left) and exclusion (lower right) tumours. Blue dots show normal exon 3 inclusion transcripts, red dots shows exon 3 exclusion. Scalebar equals to 50μm. **D)** *Ctnnb1* exon 3 exclusion calculated as % of red dots (Ex2-4probe) over total number of dots (Ex2-4 probe + Ex 2-3 probe). INC denotes exon 3 inclusion tumours and EXC denotes exon3 exclusion tumours. Individual data points, mean ± SEM are indicated. *p< 0.05, **p<0.01 (Student’s t-test) **E)** Frequency of aberrant splicing in *CTNNB1* exon 3 in human cell lines and primary tumours for HCC and hepatoblastoma. Legend indicates exclusion of exon 3, deletion of exon3-exon4, coding mutations in exon 3, and *Ctnnb1* wildtype transcripts.

To corroborate this finding and characterize the spatial and quantitative expression of this *Ctnnb1* splice variant at the single cell level, we performed BaseScope assays on paraffin sections from HCC as well as FL and RD control livers (Figure 5B-D). To this end, one probe spanning the junction of exon2-exon3 and one probe for the aberrant exon2-exon4 junction was used (Figure 5B). As expected, the exon2-3 probe was detected in all tissues while no signal from the exon2-4 probe was detected in RD control tissues, FL or tumours previously identified by RNA-seq as not having the exon 3 exclusion transcript. In contrast, in tumours characterized by sequencing as exon 3 exclusion tumours, an average of 12% of the total number of *Ctnnb1* transcripts detected by BaseScope did not include exon3, corroborating on tissue sections the results from the splicing analysis (Figure 5D). These transcripts were detected in most of the *Ctnnb1*-expressing cells throughout the tissue and co-expressed with the normal exon3 inclusion transcripts.

In addition, analysis of published RNA-seq datasets for human primary liver tumours and cell lines showed that >1% (6 out of 471 primary tumours, 1 out of 57 HCC cell lines) of HCC patients carried *Ctnnb1* transcripts excluding exon 3 (Figure 5E). Importantly, to reduce the contribution of somatic mutations to aberrant exon 3 exclusion, transcripts were only considered in this analysis if the entire exon was excluded and the whole exons 2 and 4 were present. We also detected exon 3 coding mutations in ~17% of HCC samples and genomic deletion of a large part of exon 3 together with exon 4 in ~9% of samples (Figure 5E). In none of these cases did we detect a combination of exon 3 exclusion and other types of rearrangements in this exon. This was also evident in hepatoblastoma, where exclusion of exon 3 occurred in >22% of the cases (10 out of 44 tumours, 2 out of 8 cell lines), mutations in exon 3 in >31% (14 out of 44 tumours, 4 out of 8 cell lines), and exon3-exon4 deletion only in one cell line (HepG2 cells). Moreover, combined analysis of published RNA-seq and whole genome sequencing data demonstrated that in MHCC97 cells, the exclusion of exon 3 is not accompanied by genomic mutations in the locus. This data suggests that similar to what we observed in DIAMOND mice, splicing of the *Ctnnb1* exon 3 can be disrupted in humans even in the absence of any changes in the DNA sequence. This highlights a previously unappreciated direct role of aberrant splicing as a potential molecular driver underlying oncogenic β-catenin signalling in specific cellular contexts, such as in the case of specific diets or overnutrition.

## Discussion

In this study, we analysed the transcriptional and regulatory landscape in DIAMOND mice and found increased activation together with rewiring of the Wnt gene regulatory network. This rewiring includes a switch in the expression of TCF/LEF effectors, where tumours show upregulation of *Lef1* and *Tcf7*, and downregulation of *Tcf7l1*. This switch may be triggered by the increased β-catenin levels/activity, as similar observations were made when modulating active levels of β-catenin in mammalian nephron progenitor cells [33]. The change in Wnt effectors is likely an important driver of tumour-specific gene expression, as TCF/LEF binding motifs are enriched in tumour-specific active chromatin, and given that Lef1/Tcf7 are mostly transcriptional activators while Tcf7l1 is known to have mainly repressive functions.

Numerous studies have highlighted the importance of Wnt/β-catenin-signalling in HCC tumorigenesis and mutations in components of this pathway have been described as key events for liver tumorigenesis in specific contexts [34]. It is well known that *Ctnnb1* exon 3 is a hotspot for oncogenic coding mutations [5,6,35] that lead to its constitutive activation due to disabling of phosphorylation sites required for cytoplasmic degradation of the protein. Recent studies have also exploited this feature and showed that *in vivo* CRISPR-targeting of exon3 in livers is sufficient to induce Wnt-dependent HCC [7,8]. However, to the best of our knowledge, it has not been previously reported that alternative splicing, independent of somatic mutations in the locus, can cause exclusion of *Ctnnb1* exon3 and consequent constitutive activation of the protein. In this study, we demonstrate that western diet in DIAMOND mice induces such exclusion of *Ctnnb1* exon 3 in a large subset of the tumours, and that this is not accompanied by mutations in the region of this exon. Whether a similar connection between metabolic input and exon 3 splicing exists in human patients remain to be investigated. Still, combining publicly available whole-genome and transcriptome sequencing data, we demonstrated that MHCC97 cells present *Ctnnb1* exon 3 exclusion independently of somatic mutations in the locus, thus providing a proof-of concept for the presence of this phenomenon also in human HCC.

Notably, all sequenced DIAMOND tumours show a consistent upregulation of *Ctnnb1* and *Lef1*, while *Tcf7l2* is not differentially regulated. This expression pattern of these genes is similar to human HCC subtype G6 [36,37] and contrasts somewhat with tumours generated via targeted genomic deletions of *Ctnnb1* exon3 [7] that give rise to two distinct tumour types with either all three genes upregulated or all three genes unchanged. This indicates that metabolic input may be an important modulator of the transcriptional response to aberrant Wnt-activation that should be considered in the clinical setting.

Importantly, changes in splicing patterns are also induced in FL tissue in DIAMOND mice, in line with recent reports on altered splicing in MAFLD [30–32,38]. In combination with the recent data showing that induction of β-catenin activity is sufficient to trigger liver tumorigenesis in mice [7,8], this suggests that altered splicing is a key event that promotes formation of a cellular environment that facilitates malignant transformation in this model. Changes in the expression of genes encoding specific splice factors have been reported to drive tumorigenesis or predispose to malignancy in HCC [15,16,39]. Since the Wnt-signalling pathway is overrepresented among the differential ASE in both fatty liver tissue and tumour tissue in DIAMOND mice, it raises the question of whether this is related to metabolic dysregulation of specific splice factors important for this pathway or whether it reflects a more stochastic process where enrichment of Wnt-signalling is a consequence of selection at the cellular level.

Taken together, we show that western diet induces genome-wide chromatin remodelling and gene expression changes, as well as altered splicing patterns in DIAMOND livers. These genome-wide changes contribute to the rewiring of the Wnt/β-catenin gene regulatory network that plays a pivotal role in HCC development. In particular, western diet in this model induces aberrant splicing that leads to the exclusion of *Ctnnb1* exon 3 in a subset of DIAMOND tumours. This demonstrates a new splice-dependent mechanism underlying oncogenic activation of β-catenin that is independent on somatic mutations in the locus.

Given the connection between diet and alternative splicing, new questions arise on how splicing patterns and expression of specific splice factors relate to not only caloric intake but also macronutrient composition, and how different types of diet may induce or alleviate such aberrations in different genetic backgrounds.

## Methods

### Animals and diets

All experiments were performed in compliance with national and institutional laws and are reported in accordance with the ARRIVE guidelines. The study was approved by the Regional Ethics Committee at the Court of Appeal of Northern Norrland (Ethical approval ID A39-2018). Male B6/129 mice were purchased from Sanyal Biotechnology and housed at 12:12 h light/dark cycle in a temperature/humidity controlled (22°C and 50% humidity) room and *ad libitum* feeding. Diet intervention was performed as previously described [19]. At 8 weeks of age, DIAMOND mice (n=20) were fed western diet (high-fat, high carbohydrate, Harlan TD.88137) supplemented with fructose/glucose water (23.1 g/L d-fructose + 18.9 g/L d-glucose) while littermate controls (n=10) were fed a standard chow diet (Harlan TD.7012) with normal water. After ~52 weeks mice were sacrificed. Food intake and body weight were measured weekly. The researchers were aware of which experimental groups the animals belonged to.

### Metabolic measurements, body composition and histology

For Insulin Tolerance Test, non-starved mice were injected intraperitoneally with insulin (Actrapid Penfill) (1U/kg body weight). Intraperitoneal Glucose Tolerance Test (GTT) combined with Glucose Stimulated Insulin Secretion (GSIS) were performed on 6 hours fasted and sedated mice (Hypnorm (Veta Pharma)/Midazolam (Hamlenmice)) following i.p. administration of glucose (SIGMA #G7021) (1 g/kg body weight). Blood glucose and plasma insulin levels were analysed from blood samples taken at indicated time points for both procedures using a Glucometer (Ultra 2, One Touch) and the ultra-sensitive mouse insulin ELISA kit (Chrystal Chem Inc. #90080), respectively.

Blood profiling of liver function was performed using the Mammalian Liver Profile rotor on a VetScan VS2 (Abaxis, Inc.). Liver TG and cholesterol were measured using Serum Triglyceride Determination Kit (Sigma-Aldrich #TR0100) and Cholesterol / Cholesteryl Ester Quantitation Kit (MBL #JM*-*K603*-*100) according to manufacturer’s recommendations. Body composition of live mice was measured using EchoMRI (EchoMRI LLC; EchoMRI 3-in-1). Liver and tumour histology was assessed using haematoxylin and eosin (H&E) and picrosirius red in paraffin-embedded sections using standard procedures.

### RNA-seq

#### Library preparation

Approximately 30 mg of frozen liver and tumour tissues were used for total RNA extraction using the RNeasy Plus Universal minikit (Qiagen, #73404). Integrity and quality of RNA were determined on the Bioanalyzer (Agilent, #5067-1511). 1 ug of total RNA was used for library preparation, firstly the RNA was enriched for mRNA using the NEBNext Poly(A) mRNA Magnetic Isolation Module (NEB, E7490) followed by the NEBNext Ultra II RNA Library Prep Kit for Illumina (NEB, E7770). The quality and concentration of libraries were assessed with Bioanalyzer (Agilent, #5067-4626) and Qubit (Thermofisher, #Q33230).

150 bp paired-end sequences were obtained using the NovaSeq 6000 Illumina Sequencer, with a coverage of ~ 37 million reads per library. Quality control of fast files was done with FastQC.

#### Gene expression analysis

Raw reads were aligned to the mouse genome (mm10, UCSC) using STAR (options: -- outSAMtype BAM SortedByCoordinate --seedSearchStartLmax 12 -- outFilterScoreMinOverLread 0.3 --alignSJoverhangMin 15 --outFilterMismatchNmax 33 -- outFilterMatchNminOverLread 0 --outFilterType BySJout --outSAMattributes NH HI AS NM MD --outSAMstrandField intronMotif --quantMode GeneCounts) [40].

Genes with a minimum of 10 reads along all the samples were kept for further analysis. Normalisation and differential expression analysis were performed using DESeq2 (Love, 2014). Genes were considered differentially expressed with pvalue < 0.01 and FDR < 0.01 (False Discovery Rate). The differentially expressed genes were grouped into early, gradual, tumour-specific, fatty liver-specific, and switching genes. Early genes were defined as genes differentially expressed in both HCC and FL compared to RD. Gradual genes included differentially expressed genes in FL vs RD but that are also increasingly dysregulated in HCC vs FL, as well as genes differentially expressed in HCC vs RD but not compared to FL. Tumour-specific genes were differentially expressed in tumour samples compared to both RD and FL while fatty liver-specific genes were differentially expressed in FL samples compared to RD, HCC or both. Lastly, a small group of switching-genes were identified that were differentially up or down in FL samples compared to RD, and the opposite in HCC samples. Gene Ontology (GO) and KEGG enrichment analysis for each cluster was performed using clusterProfiler (pvalue < 0.01 and FDR < 0.01) [41]. For pairwise comparisons of gene expression of selected genes in the Wnt-signalling pathway, normalized counts and the adjusted p-value obtained with DESeq2 was used.

#### Alternative splicing analysis

PSI values were calculated using vast-tools with default parameters [42]. An event was considered an ASE in a sample if its PSI value is between 10% and 90% and the read coverage was sufficient (minimum of “VLOW” coverage score or higher in vast-tools). A t-test was performed to compare the total number of events, total number of genes with ASE or the ratio of ASE/gene per group. RNA-seq datasets of human MAFLD samples (control: n = 10 and MAFLD: n = 206) were obtained from GSE135251 [43]. Statistical significance on comparisons of *Ctnnb1* exon 3 exclusion between the groups was calculated with vast-tool default parameters.

An event was considered differentially spliced if the absolute estimated dPSI was higher than 10% and higher than the minimum value of dPSI at a 99% confidence based on vast-tools diff software [42].

### ChlP-seq

#### Library preparation

For ChIP-seq experiments approximately 200 mg of fresh liver and tumour tissue was cut into smaller pieces and fixed for 15 min in 1 ml fixing solution (1% formaldehyde, 50 mM HEPES–KOH, 100 mM NaCl, 1 mM ethylenediaminetetraacetic acid (EDTA), 0.5 mM EGTA) at RT with gentle mixing. Cross-linking was stopped by adding 1/20 volume of 2.5 M glycine for 5 min at RT. Samples were then washed 2 × 5 min at 4 °C with cold 1 × PBS containing protease inhibitor cocktail (Roche, #4693132001), snap frozen and stored at −80°C until awaiting further processing. Liver and tumour tissues were pulverised before lysis on cell lysis buffer at 4°C for 5-10 minutes. Nuclei were pelleted and resuspended in 1xPBS and the volume corresponding to 50-70 mg of tissue was used for chromatin extraction by adding nuclei lysis buffer at 4°C for 5 minutes. Chromatin was sonicated on a Covaris E220 device (PIP: 105, Duty factor: 2, Cycles/burst: 200, Duration: 5 min). ChIP was performed with anti-H3K27ac (Abcam, ab4729) and anti-H3K27me3 (Abcam, ab192985) antibodies. ChIP-seq libraries were prepared with the NEBNext Ultra II DNA Library Prep Kit for Illumina (NEB, E7645) and quality and concentration were assessed with the Bioanalyzer (Agilent, #5067-4626) and Qubit (Thermofisher, #Q33230).

150 bp paired-end sequences were obtained using the NovaSeq 6000 Illumina Sequencer, with coverage of ~ 52 million reads per library. Quality control of fast files was done with FastQC.

#### Data analysis

Raw reads were mapped to the mouse genome (mm10, UCSC) using bowtie2 (options: -- threads 4 —very-sensitive) [44]. Samtools was used to convert SAM files into BAM files. Enriched peaks were called using MACS2 (options: callpeak --gsize 1.87e9 --broad --broad-cutoff 0.05 --f BAMPE) [45]. DiffBind package was used for differential binding analysis [46,47]. Peaks were annotated using ChIPseeker [48] and Gene Ontology (GO) and KEGG enrichment analysis for each cluster was performed using clusterProfiler (pvalue < 0.01 and FDR < 0.01) [41]. Motif enrichment analysis for each subset of regions was done using HOMER with default settings [49].

### *Ctnnb1* exon 3 exclusion in datasets from human subjects

Previously published human HCC and hepatoblastoma datasets were obtained from GSE97098 [50], GSE193567 [51], GSE184733 [52], GSE65485 [53], PRJNA867011 [54], PRJNA870935 (https://www.ncbi.nlm.nih.gov/bioproject/?term=PRJNA870935), PRJEB21899 [55], GSE183406 [56], GSE83518 [57], and GSE133039 [58].

Raw reads were aligned to the human genome (hg38, UCSC) using STAR (options: -- outSAMtype BAM SortedByCoordinate --seedSearchStartLmax 12 --outFilterScoreMinOverLread 0.3 --alignSJoverhangMin 15 --outFilterMismatchNmax 33 -- outFilterMatchNminOverLread 0 --outFilterType BySJout --outSAMattributes NH HI AS NM MD --outSAMstrandField intronMotif --quantMode GeneCounts) [40]. Bam files were visualised in IGV to find samples with *Ctnnb1* exon skipping or mutations.

### RT-PCR and genomic PCR

*Ctnnb1* exon 3 skipping was confirmed in several tumour samples by extracting total RNA from 5-20 mg of tissue using the RNeasy Plus Universal minikit (Qiagen, #73404). cDNA was synthesized using the RevertAid protocol (ThermoScientific, EP0451).

Genomic DNA was extracted by incubating 5 mg of tissue in 50mM NaOH for 25 minutes at 95°C and then 1/10th of 1M Tris-HCl pH 8. Primers flanking the exon3 were used for PCR amplification on both the cDNA and genomic DNA to amplify the region (*Ctnnb1* exon2 Fw: GGTACCTGAAGCTCAGCGCA and *Ctnnb1* exon4 Rv: CTGGTCCTCATCGTTTAGCA). Gel bands were purified using the NucleoSpin Gel and PCR Clean-up kit (Macherey-Nagel, N. 740609.50) and sequenced by Sanger sequencing.

### BaseScope RNA In Situ Hybridisation (ISH)

We employed the BaseScope RNA ISH assay (ACD, cat. no 323871) to detect the *Ctnnb1* exon3 skipped isoform. In brief, we utilised custom-designed probes from Advanced Cell Diagnostic (ACD) (Newark, CA, USA): (1) a mouse-specific probe (1ZZ) (BA-Mm-Ctnnb1-1zz-st-C1) that targets the exon2-exon3 junction to detect the normal *Ctnnb1* and (2) a mouse-specific probe (1zz) (BA-Mm-Ctnnb1-E2E4-C2) that targets the exon2-exon4 junction to detect the novel *Ctnnb1* exon3 skipped isoform. Each batch run of the assay included an RD, an FL, an HCC exon3 included and an HCC exon3 excluded (as determined by RT-PCR) sample.

Slides were scanned on a Zeiss Axioscan and quantification was done manually assisted by the Qupath software. The number of dots per field was determined in a total of 10 fields per sample. Data was represented as % of exon3 exclusion which was calculated as percentual ratio of dots from exon2-4 probe vs total (exon2-4 / (exon2-4 + exon2-3 dots). Statistical significance was assessed using a two-tailed Student’s t-test.

### Statistical analysis

Prism 9 (GraphPad Software, LLC) was used for statistical analysis of body composition and metabolic parameters. Statistical methods for all analysis are indicated in figure legends and in the methods section for each experiment. Normality and log-normality tests were performed to select the appropriate statistical methods for analysis. RNA-seq and ChIP-seq data were analysed using specific R-packages (see separate section).

## Data availability statement

All generated datasets for RNA-seq and ChIP-seq are available through the Gene Expression Omnibus repository under the accession number GSE250577. Additional data that support the finding of this study are available on reasonable request from the corresponding author.

## Acknowledgements

We thank Ingela Lundberg for animal care and members of the Edlund laboratory for technical assistance on metabolic experiments. We also acknowledge the facilities and technical assistance of Umeå Center for Comparative Biology (UCCB). The computations were enabled by resources in project SNIC 2021/23-439, 2321/22-541, 2022/23-376 and 2022/22-671 provided by the Swedish National Infrastructure for Computing (SNIC) at UPPMAX, partially funded by the Swedish Research Council through grant agreement no. 2018-05973. Finally, we acknowledge past and present members of the Hörnblad and Remeseiro laboratories for helpful discussions.

