## Supplemental data for "Diet-induced rewiring of the Wnt gene regulatory network connects aberrant splicing to fatty liver and liver cancer in DIAMOND mice"

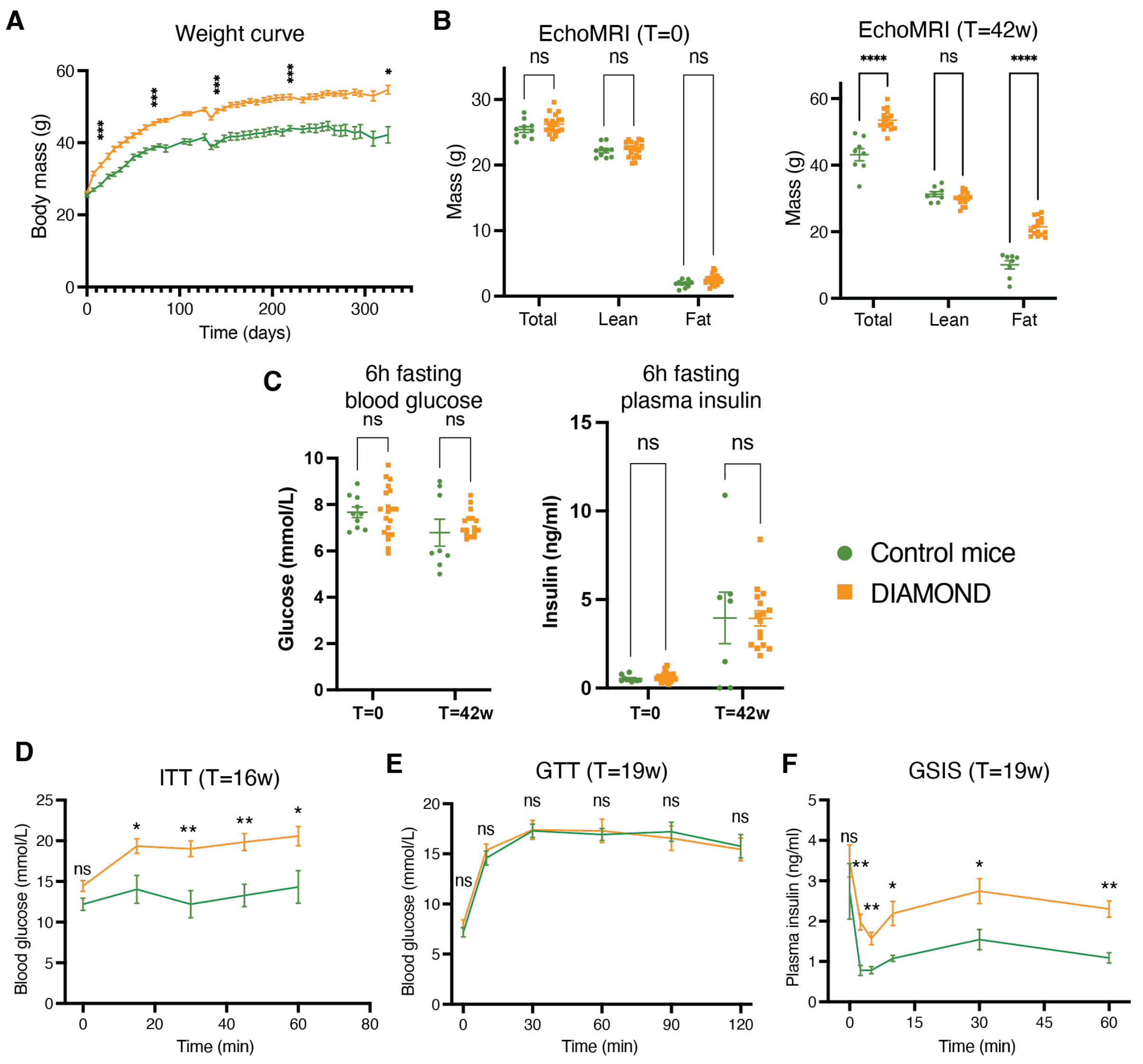


**Figure S1. Obesity and insulin resistance in DIAMOND mice. A**) Body weight change over time in mice fed RD (Control mice, green line) and mice fed WD (DIAMOND mice, orange line). Mean ± SEM are indicated. Difference in body mass is statistically significant at all time points from T=7days. For simplification only 5 time points are depicted in the graph. *p< 0.05, ***p<0.001. **B**) Body mass composition, **C**) fasted blood glucose (left), and fasted insulin (right) at start of the experiment (T=0) and after 42 weeks of diet (T=42w) for controls (green circles) and DIAMOND (orange squares). Individual data points, mean ± SEM are indicated. ****p<0.0001, ns = not significant (Student´s t-test). Blood glucose levels during **D**) Insulin tolerance test (ITT), **E**) Glucose Tolerance Test (GTT), and **F**) insulin plasma profile during Glucose-Stimulate Insulin Secretion test (GSIS) for controls (green) and DIAMOND (orange). Mean ± SEM are indicated. *adj.p< 0.05, **adj.p<0.01, ns = not significant (Student´s t-test).

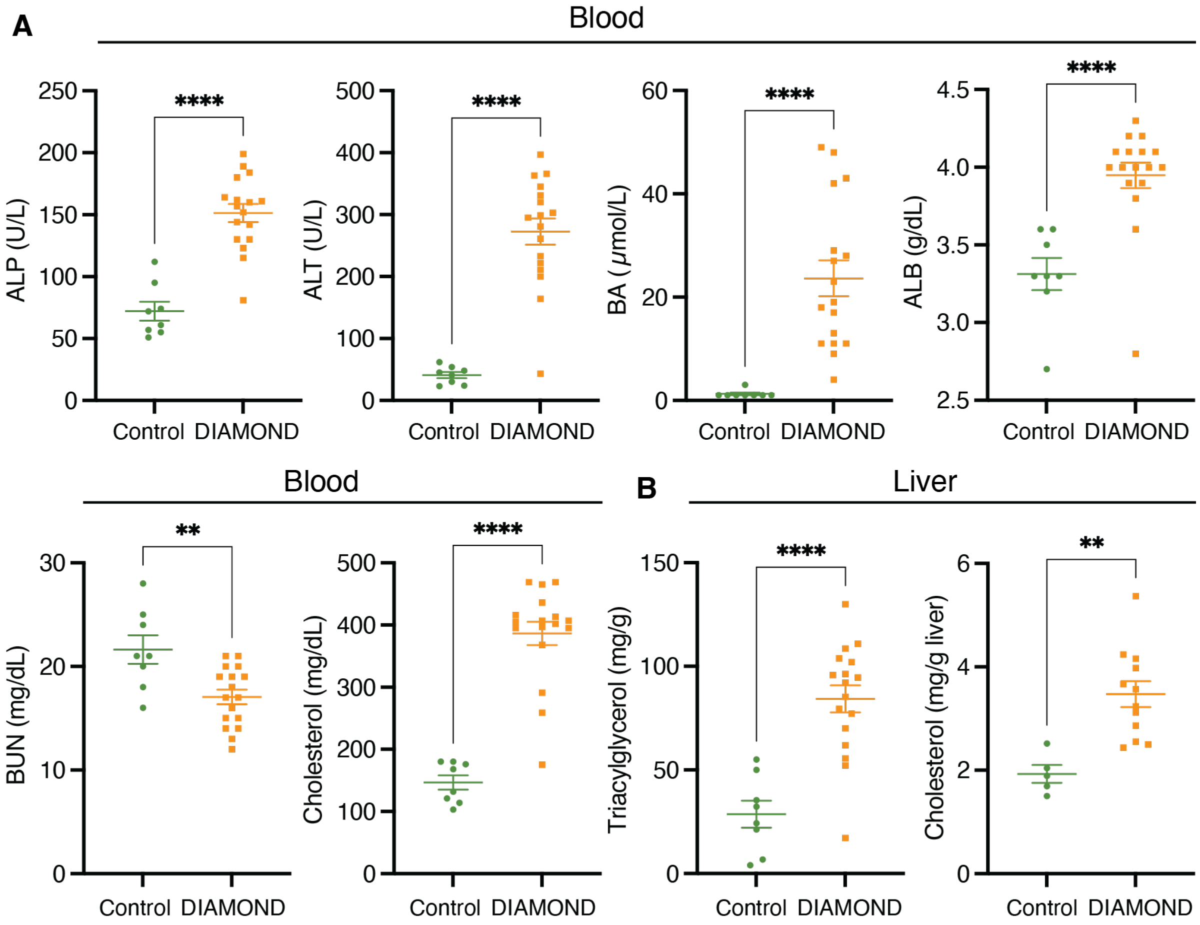


**Figure S2. Disturbed liver function and dyslipidemia in DIAMOND mice. A**) Blood levels of alkaline phosphatase (ALP), aspartate transaminase (ALT), bile acids (BA), albumin (ALB), urea nitrogen (BUN) and cholesterol at T=38w for mice fed regular diet (controls, green) and western diet (DIAMOND, orange). Individual data points, mean ± SEM are indicated. **p<0.01, ****p<0.0001 (Student´s t-test for ALP, ALT and BUN, Mann-Whitney test for BA, ALB, and cholesterol). **B**) Liver triacylglycerol and cholesterol levels in the same mice (controls: green, DIAMOND: orange) **p<0.01, ****p<0.0001 (Student´s t-test).

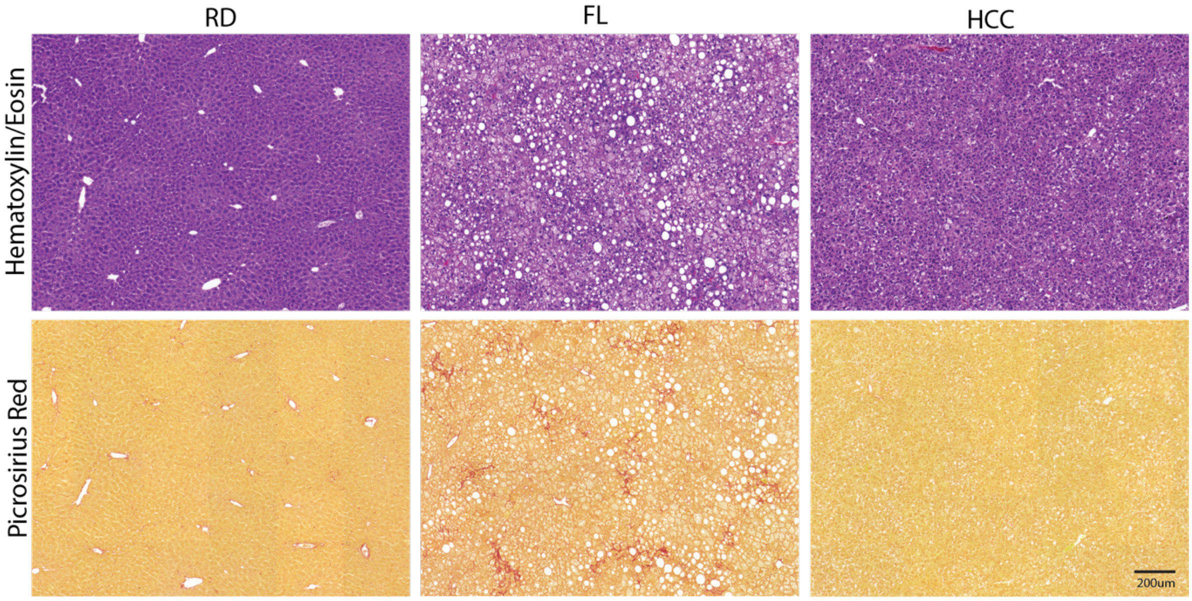


**Figure S3. Histology of DIAMOND liver tissue and tumours.** Representative photomicrographs of sections from DIAMOND RD livers, FL tissue, and HCC tumours stained for hematoxylin and eosin (upper row) and picrosirius red (lower row). In contrast to fatty liver tissue, the tumours are devoid of fibrosis and lipid accumulation.

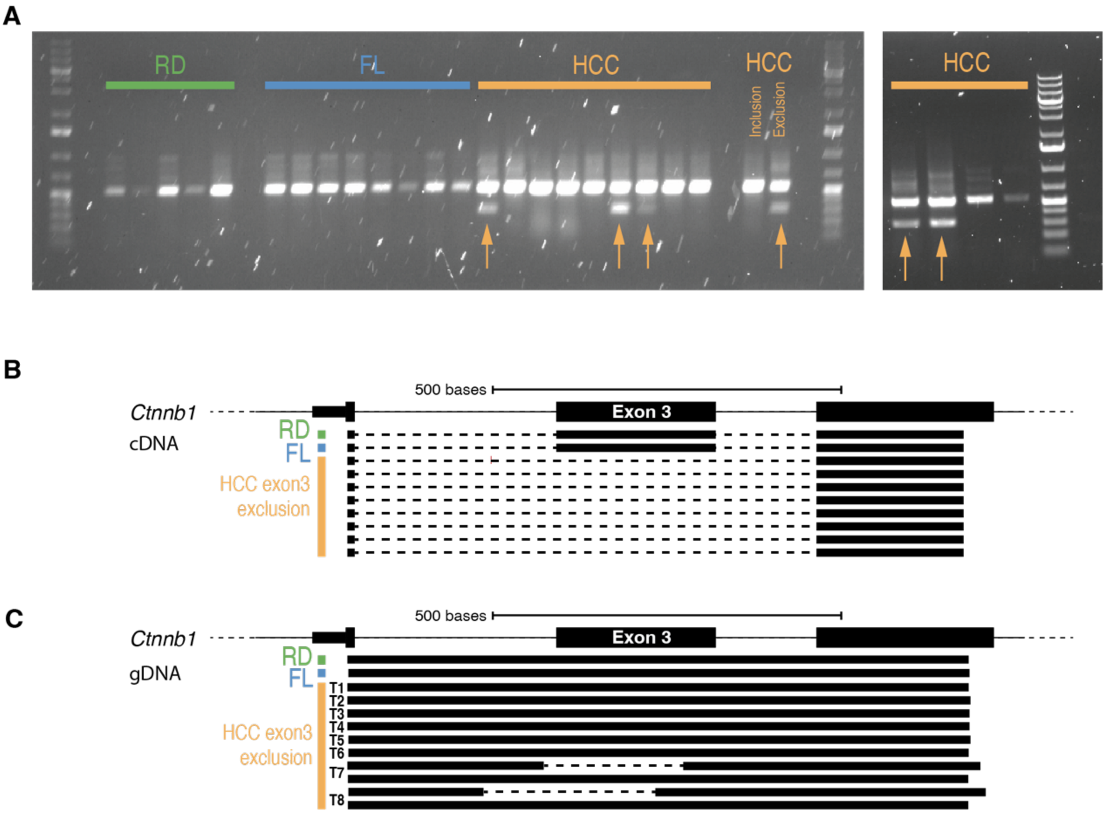


**Figure S4. DIAMOND tumours display Ctnnb1 Exon3 exclusion independent on genomic mutations in the region. A**) Agarose gel electrophoresis of PCR amplified cDNA from RD liver, FL and HCC tissue. Arrows indicate exon 3 exclusion amplicon. One tumour not excluding exon 3 ("Inclusion") and one tumour with exon 3 exclusion transcripts ("Exclusion") from RNA-seq data were used as controls. **B**) Alignment of Sanger-sequenced Ctnnb1 transcripts to genomic region. Normal Ctnnb1 transcripts for 1 RD control liver (green) and 1 FL tissue (blue) are displayed in upper two rows, and Exon 3 exclusion transcripts (orange) are displayed below. **C**) Alignment of genomic sequence in the Ctnnb1 exon 3 region for 1 RD liver (green), 1 FL tissue (blue) and the Ctnnb1 Exon 3 exclusion tumour tissue (orange) from **B**). Note that only two out of 8 exon 3 exclusion tumours have genomic deletions in the region (T7 and T8). For these, both the wild type allele and the deletion are depicted.
